# Early disruptions in vitamin D receptor signaling induces persistent developmental behavior deficits in zebrafish larvae

**DOI:** 10.1101/2025.03.28.645997

**Authors:** Morgan Barnes, Derek Burton, Kurt Marsden, Seth Kullman

## Abstract

A critical function of the nervous system is to rapidly process sensory information and initiate appropriate behavioral responses. Defects in sensory processing and behavior selection are commonly observed in neuro-psychiatric conditions including anxiety, autism (ASD), and schizophrenia. The etiology of sensory processing disorders remains equivocal; however, it is hypothesized that extrinsic environmental factors can play fundamental roles. In this study we examine the importance of vitamin D (1α, 25-dihydroxyvitamin D3) receptor signaling during early life stage development on sensory processing and neurobehavioral health outcomes. While vitamin D has traditionally been associated with mineral ion homeostasis, accumulating evidence suggests non-calcemic roles for vitamin D including early neurodevelopment. Here we demonstrate that systemic disruption of vitamin D receptor (VDR) signaling with a conditional dominant negative (dnVDR) transgenic zebrafish line results in specific visual and acoustic sensorimotor behavior defects. Induction of dnVDR between 24-72 hours post fertilization (hpf) results in modulation of visual motor response with demonstrate attenuation in acute activity and hypolocomotion across multiple swimming metrics when assayed at 6- and 28-days post fertilization (dpf). Disruption in VDR signaling additionally resulted in a strong and specific attenuation of the Long-Latency C-bends (LLC) within the acoustic startle response at 6 dpf while Short-Latency C-bends (SLC) were moderately impacted. Pre-pulse inhibition (PPI) was not impacted in young larvae however exhibited a significantly attenuated response at 28 dpf suggesting an inability to properly modulate their startle responses later in development and persistent effects of VDR modulation during early development. Overall, our data demonstrate that modulation of vitamin D signaling during critical windows of development irreversibly disrupts the development of neuronal circuitry associated with sensory processing behaviors which may have significant implications to neurobehavioral health outcomes.

## Introduction

Neuropsychiatric disorders such as anxiety, autism (ASD), and schizophrenia (SZ) typically have defects in sensory processing and behavior[1]. The nervous system is critical for processing sensory information and initiating appropriate behaviors. Despite the biological and clinical relevance, our understanding of the cellular and molecular mechanisms regulating these processes is limited, in part due to the intricate and dynamic circuitry involved in complex human decision-making. The origins of sensory processing disorders are complex, as intrinsic/genetic, extrinsic/environmental and the interactions of intrinsic and extrinsic factors can play fundamental roles[2]. Low levels of vitamin D during early stages of development have been linked to various brain disorders such as schizophrenia and autism with associated deficits in sensory processing[3]. However, to date, there remain critical gaps in the knowledge of how vitamin D deficiency (VDD) contributes to neurobehavioral health outcomes.

Vitamin D was first identified with the investigation of rickets by McCollum who coined the term vitamin D[4]. Subsequently it was demonstrated that vitamin D was essential for dietary calcium absorption and skeletal mineralization[5]. Research has since illustrated that vitamin D serves many physiological roles and vitamin D deficiency is now associated with select disease etiologies including cardiovascular disease, cancer, stroke, metabolic disorders and neurodegenerative disease[6, 7].

Vitamin D is a fat-soluble vitamin that can be obtained through the diet or UVB conversion of 7-dehydrocholesterol to pre-vitamin D3 in the skin followed by metabolic activation to vitamin D3. There are multiple forms that circulate throughout the body including the inactive 25 hydroxyvitamin D, (25(OH)D) and the active form 1,25-hydroxyvitamin D (1,25(OH)2D). 1,25-hydroxyvitamin D is formed through sequential hydroxylations involving both 25-hydroxylase and 1α-hydroxylase[8]. The active form of vitamin D serves as a potent ligand for the vitamin D receptor (VDR) and together with its obligate heterodimerization partner RXR, this receptor pair facilitates both transactivation and trans-repression of genes critical to select cellular processes including maintenance of calcium and phosphate homeostasis, cell proliferation, immune system function, cardiovascular function, and neurodevelopment[8].

Vitamin D deficiency is defined as 1,25-hydroxyvitamin D levels lower than 50 nmol/L or 20 ng/mL and multiple factors can affect vitamin D levels including age, lifestyle, skin pigmentation, exposure to environmental agents, diseases that affect the intestinal tract and people who have undergone gastric bypass surgery[8, 9]. About a billion people worldwide are vitamin D deficient (VDD) or insufficient, making this a global issue[10]. The prevalence of VDD in pregnant women, who are especially vulnerable, can be as high as 50% globally if not more[11]. Pregnant women are at particularly high risk for VDD, due to increased demand for vitamin D to supplement the growing fetus, which is associated with indirect and direct adverse neurodevelopmental effects[11]. The vitamin D receptor (VDR) itself has been found to be expressed within multiple regions of the mammalian brain in conjunction with 1α-hydroxylase, an essential enzyme necessary to convert inactive vitamin D to the active form[12]. Multiple studies indicate a pivotal role for vitamin D as a neurosteroid in the brain mediating essential neural functions including regulation of calcium homeostasis[13], deposition of beta-amyloid[14, 15], regulation of oxidative stress[16, 17], inflammation[17, 18], cell proliferation[19, 20], cell differentiation[19, 21], regulation of neurotropic factors[22–25] and biosynthesis of neurotransmitters[26–29]. These actions have categorized vitamin D as a neuroprotectant and has identified vitamin D as being critical to brain health[3, 6].

Translational models including zebrafish (*Danio rerio*) have proved a valuable resource to investigate developmental neuroscience research. The zebrafish genome is well annotated, and it has been demonstrated that zebrafish share about 70% genetic homology to humans[30]. Additionally, zebrafish possess similar neurochemistry to humans including neurotransmitter receptors, transporters and synthesizing enzymes[30]. The zebrafish central nervous system (CNS) begins to be specified as early as 6 hours post fertilization (hpf), at the beginning of gastrulation[30]. At approximately one week old, zebrafish brains are estimated to contain roughly 100,000 neurons and can perform many complex behaviors despite being so young[31]. Zebrafish have been demonstrated to be similar to mammalian research models via brain macro-organization and cellular morphology[32]. CNS structures in zebrafish can be identified as early as 10 hpf, and by 24 hpf the primary divisions of the brain, the forebrain, midbrain and hindbrain, are defined by morphogenetic boundaries. Additionally, zebrafish larvae possess most of the major neurotransmitter’s receptors, transporters and enzymes[32]. Zebrafish as young as 2 days post fertilization (dpf) are able to respond to external stimuli through sensorimotor integration in the CNS[30]. Zebrafish have a very social nature and are diurnal animals that rely on vision, similar to most humans[32]. Over the past several decades multiple assays have been developed to index a broad range of zebrafish behaviors including scoot swimming; burst swimming; routine turns; stimulus-specific turning responses such as J-, C-, and O-bend turns; optokinetic and optomotor responses; prey tracking; phototaxis; thigmotaxis; escape and avoidance behaviors; non-associative learning; and visual recognition memory[33, 34]. These assays have proven to be sensitive to the early and persisting impacts of exogenous drug/toxicant exposures, dietary modifications and/or genetic manipulations[35, 36]. These tests provide a powerful approach to assess alterations within early neurodevelopment and sensory processing, with defects reflecting dysfunction of the underlying cellularity and/or neural circuits.

In this study we take advantage of a set of well-established behavioral assays to investigate how modulation of developmental VDR signaling modulates larval zebrafish sensory processing. Specifically, we use an inducible dominant-negative VDR transgenic line to simulate VDD by limiting embryonic/larval VDR signaling, and we show that VDR signaling is critical within a defined developmental window for normal responses to both visual and auditory stimuli. Furthermore, we find that some of these behavioral changes persist into young adulthood, and they are consistent with changes in the expression of genes in several key neural signaling pathways.

## Methods

### Fish Husbandry and Maintenance

Zebrafish (Danio rerio) were housed and cared for according to standard protocols approved by the North Carolina State University (NC State) Institutional Animal Care and Use Committee. Adult zebrafish were maintained at appropriate densities in 9L tanks as part of a recirculating aquatics system under a 14:10hr light: dark cycle. Water temperature was maintained at 28.5±0.5°C with a pH between 6.8 and 7.5. Two strains of zebrafish (Danio rerio) were used in this study including the wildtype strain EK and a heat shock inducible vitamin D receptor dominant negative transgenic strain TG(hsp:dnVDRa:BFP) obtained from the Poss lab, Duke University[37]. For embryo collection, mature zebrafish (4-12 months) were bred under standard conditions and viable embryos were sorted and collected. Embryos were maintained in petri dishes with E3 media in an incubator set to 28 ± 0.5 °C. Embryo media was changed daily.

### Heat Shock Induction

Heat shock was performed on zebrafish embryos at either 24, 48 or 72 hours post fertilization (hpf) using a thermocycler. Heat shock was induced at indicated times by placing embryos/larvae in a 96 well plate, 1 fish per well, with E3 media then incubated in a thermocycler at 38°C for 30 minutes. Embryos/larvae were subsequently screened 5 to 8 hours post heat shock for induction of blue fluorescent protein (BFP) expression using a UV filter on an epi-fluorescent (Nikon Eclipse TE2000-S) microscope, and fish were sorted into two cohorts according to the presence (+) or absence (-) of BFP expression which is linked to dnVDR with a P2A sequence.

### Acoustic Startle Response

Vibro-acoustic startle responses were assayed using custom built behavioral systems at NC State. We exposed 6 dpf larvae to 10 trials at each of 6 intensities of acoustic stimuli (ranging from 10-60 dB) at a 20 s inter-stimulus interval (ISI) for a total of 60 stimuli. To assess prepulse inhibition (PPI), larvae were exposed to 10 pairs of a weak (23 dB) prepulse stimuli and strong (60 dB) stimuli. There was a 300 ms break between the prepulse and pulse, and a 20 s ISI between pairs of stimuli. For 28 dpf fish, which habituate more readily than younger fish, we ran 5 trials at each of 3 intensities, with a 2 min ISI to assess startle responsiveness, followed by 10 PPI trials. We captured responses with a Photron Mini-UX50 high-speed camera at 1000 frames/s. To evaluate movement kinematics (response frequency, latency, turning angle) we used the FLOTE software package[38]. For each phase of the assay, we measured multiple endpoints, including response frequency, latency, turning angles, angular velocity, duration and distance traveled. A minimum of 20 larvae were tested for each replicate set, with three replicate analyses performed.

### Zebrafish Larval visuo-motor response (VMR) assay

The larval VMR behavioral analysis was conducted using the Noldus video-surveillance system and animal movement tracking software EthoVision XT (Noldus Information Technology, The Netherlands). All larval testing was run between 9:00 AM and 1:00 PM to ensure consistency with circadian cycles. Larval locomotion assays were conducted using a DanioVision™ lightbox running EthoVision XT® tracking software (Noldus Inc., Wageningen, The Netherlands). Heat shocked, 6-day or 28-day old larvae were transferred to multi-well plates, with all experimental groups represented on each plate and across multiple plates. Following a 1-hour incubation at 28°C, locomotor activity is tracked during an initial 30-min acclimation period in the dark (0% illumination), followed by five cycles of 10 min at 100% illumination (5000 lx) and 10 min at 0% illumination. An infrared camera captures larval locomotion across the programmed cycles. For each trial the average distance moved, total velocity, activity state, and distance to point is determined for each subject. The raw data of movement per minute for the alternating 10-min dark and light phases of the session was utilized for the statistical analyses and a minimum of 20 larvae was tested for each replicate set, with three replicate analyses performed.

### Epifluorescent imaging of BFP expression

To localize BFP expression post heat shock tg(hsp:dnVDRa:p2a:BFP) zebrafish were anesthetized using 0.006% tricaine then placed on a slide. Images were collected using Elements Software and camera.

### RNA Isolation

Larval zebrafish (6 dpf) were euthanized in ice water. 15 to 20 whole larvae we subsequently pooled per RNA sample with 4 replicated samples. Samples were immediately flash frozen in liquid nitrogen and tissues were homogenized in TRI Reagent® (Ambion®, Life Technologies, Carlsbad, CA, USA) using a handheld BioVortexer (Thomas Scientific, Swedesboro, NJ, USA). Total RNA was isolated according to the TRI Reagent manufacturer’s protocol. Total RNA was quantified using the Qubit® RNA HS Assay Kit (Invitrogen™, USA) containing Qubit® RNA HS Reagent and Buffer with the Qubit® 3.0.

### Quantitative Real-Time PCR (qPCR)

Quantitative Real-Time PCR (qPCR) was used to measure targeted gene expression in whole 6 dpf larval zebrafish. cDNA was synthesized from total RNA using 10x random primers, 10x reverse transcription buffer, MultiScribe Reverse Transcriptase, and 10mM deoxynucleotide triphosphates from a High-Capacity cDNA Reverse Transcription Kit (Applied Biosystems, Foster City, CA) along with RNasin(R) RNase Inhibitor (Promega, Madison, WI). Primer sequences were designed using Primer3web, version 4.1.0. Primers were ordered from Integrated DNA Technologies, Inc. (Coralville, Iowa). See **Supplemental Table 1** for a full list of primer sequences. Gene expression patterns were quantified using a QuantStudio 3 real time PCR machine. Biological replicates (n=3–4/group) were plated in triplicates and amplified in a 96-well, clear Olympus PCR plate (Genesee, Morrisville, NC). Each well contained a 20µL mixture of Ultrapure water (Invitrogen, Marietta, OH), forward primer, reverse primer, cDNA, and iTaq Universal SYBR Green Supermix (Bio-Rad, Hercules, CA). Each reaction occurred under the following conditions: (1) 50 °C for 2 min, (2) 95 °C for 10 min, (3) 95 °C for 15 sec followed by 60°C for 1 min (repeated 40 times). This cycle was followed by a dissociation stage which ensured primer specificity and confirmed the absence of primer dimerization: (4) 95 °C for 15 secs 60 °C for 1 min, 95 °C for 15 sec, 60 °C for 1 min. Individual threshold cycle values (Ct) were determined for each reaction by the QuantStudio 3 Software and relative fold change differences for each gene across each sample was calculated according to the ΔΔCt method[39]. Gene expression was normalized to efla as the housekeeping gene.

## Results

### Heat shock induction of dnVDR:BFP

Induction of heat shock at 24, 48, 72 hours post fertilization induced robust BFP expression in Tg:dnVDR:p2A:BFP zebrafish larvae as observed under epifluorescence. A representative image of BFP expression is shown in Supplementary Figure 1. BFP expression depicted in our cohort was comparable to BFP expression shown in previous studies with strong localization in the brain[37]. The weak peptide bonds within the p2A linker allows for BFP to mark the cells in which dnVDR is expressed without interfering with its function. Post heat shock zebrafish larvae were observed under bright field microscopy to ensure heat shock treatment did not result in any observable morphological alterations to zebrafish larvae.

### Induction of dnVDR modulates early life stage visual motor response (VMR) behavior

To assess deficits in visual motor sensory processing we conducted the light dark locomotor activity (visuomotor response, VMR) assay at 6 dpf following heat shock induction of dnVDR at either 24, 48, or 72 hpf. Results of this analysis demonstrate that induction of the dominant negative vitamin D receptor within these time periods altered behavioral outcomes across four swimming parameters including distance moved, velocity, activity state, and distance to center point of the well (**Figure 1A-D** and **Table 1A**). Induction of the dnVDRa at 24 hpf results in significant reductions in the dark phase total distance moved (p<0.0001), velocity (p<0.0001) and activity state (p<0.0001). Induction of the dnVDRa at 24 hpf also significantly reduced the light phase total distance moved (p<0.001) and velocity (p<0.05), while increasing distance to point (p<0.01). The second timepoint of dnVDRa induction, 48 hpf, caused similar changes in the dark phase for total distance moved (p<0.0001), velocity (p<0.05) and activity state (p<0.01). Induction of the dnVDRa at 48 hpf also resulted in significant alterations in the light phase in total distance moved (p<0.05) and distance to point (p<0.01). The final timepoint of dnVDRa induction, 72 hpf, resulted in the least severe effects with alterations only in the light phase in total distance moved (p<0.05) and velocity (p<0.05). These results indicate that disruption of vitamin D receptor signaling via induction of the dnVDRa at early timepoints at 24 and 48 hpf has severe impacts on light dark locomotor activity.

**Figure 1:**
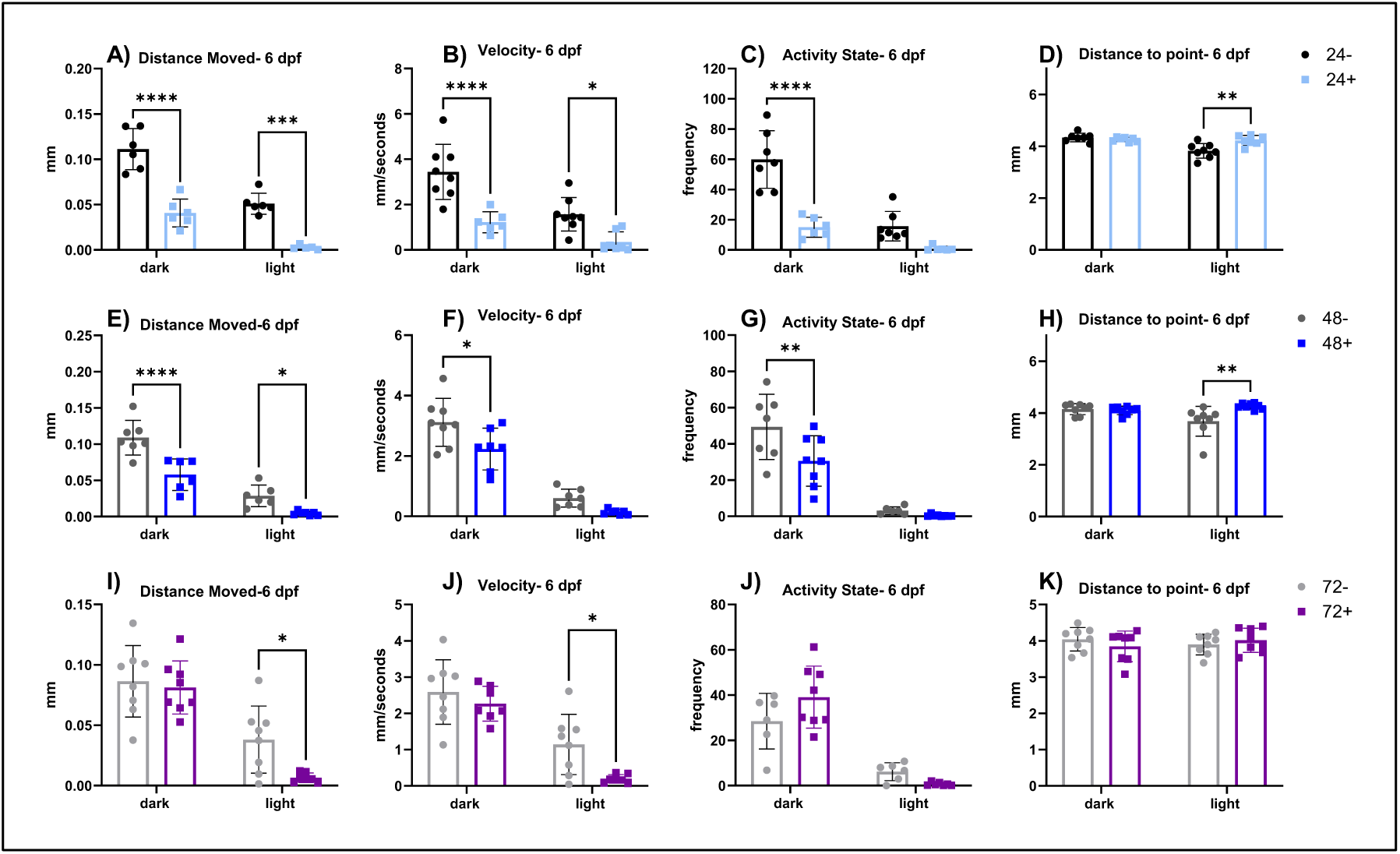
Parameters of VMR separated by light or dark periods run at 6 dpf. (A-D) VMR parameters in dnVDRa induced zebrafish at 24 hpf. There is a significant decrease in distance moved in the dark (p<0.0001) and light periods (p<0.001), a decrease in velocity in the light (p<0.0001) and dark periods (p<0.05), a decrease in activity state in the dark period (p<0.0001) and an increase in distance to point in the light period (p<0.01) in the 24+ fish. (E-H) VMR parameters in dnVDRa induced zebrafish at 48 hpf. There is a significant decrease in distance moved in the dark (p<0.0001) and light (p<0.05) periods, a significant decrease in velocity in the dark period (p<0.05), a significant decrease in activity state in the dark period (p<0.01) and a significant increase in distance to point in the light period (p<0.01) in the 48+ fish. (I-K) VMR parameters in dnVDRa induced zebrafish at 72 hpf. There is a significant decrease in distance moved (p<0.05) and a significant decrease in velocity the light period (p<0.05) in the 72+ fish in the light period.

**Table 1:** All behavioral data values at 6 dpf. (1A) Data values for all dnVDRa induced timepoints (24+, 48+ and 72+) and their controls (24-, 48-, 72-) for all VMR parameters. (1B) Data values for all dnVDRa induced timepoints (24+, 48+ and 72+) and their controls (24-, 48-, 72-) for all ASR parameters. Bolded values represent statistical difference between the dnVDRa induced fish versus their controls.

| Table 1A: Visuo-motor response (VMR) assay at 6 dpf |  |  |  |  |  |  |  |  |  |  |  |  |  |  |  |  |
| --- | --- | --- | --- | --- | --- | --- | --- | --- | --- | --- | --- | --- | --- | --- | --- | --- |
| Parameter | Distance Moved (mm) |  |  |  | Velocity (mm/s) |  |  |  | Activity State (frequency) |  |  |  | Distance Moved (mm) |  |  |  |
| Bin | Dark |  | Light |  | Dark |  | Light |  | Dark |  | Light |  | Dark |  | Light |  |
| Data | average | SD | average | SD | average | SD | average | SD | average | SD | average | SD | average | SD | average | SD |
| 24- | 0.111 | 0.021 | 0.051 | 0.011 | 3.441 | 1.136 | 1.570 | 0.690 | 59.861 | 17.651 | 15.700 | 9.015 | 4.336 | 0.153 | 3.822 | 0.263 |
| 24+ | **0.041** | **0.014** | **0.003** | **0.002** | **1.221** | **0.424** | **0.344** | **0.416** | **14.988** | **5.987** | 0.873 | 1.422 | 4.261 | 0.079 | **4.223** | **0.179** |
| 48- | 0.109 | 0.022 | 0.029 | 0.014 | 3.116 | 0.743 | 0.603 | 0.276 | 49.361 | 16.673 | 3.137 | 1.871 | 4.151 | 0.199 | 3.684 | 0.536 |
| 48+ | **0.058** | **0.020** | **0.005** | **0.003** | **2.228** | **0.639** | 0.136 | 0.078 | **30.547** | **13.049** | 0.435 | 0.538 | 4.100 | 0.152 | **4.263** | **0.104** |
| 72- | 0.086 | 0.028 | 0.038 | 0.026 | 2.590 | 0.831 | 1.143 | 0.777 | 28.454 | 11.207 | 6.190 | 3.624 | 4.043 | 0.305 | 3.898 | 0.265 |
| 72+ | 0.081 | 0.021 | **0.007** | **0.004** | 2.265 | 0.444 | **0.195** | **0.109** | 39.078 | 12.843 | 0.720 | 0.674 | 3.848 | 0.397 | 4.014 | 0.312 |

**Table 1:** All behavioral data values at 6 dpf.
| Table 1B: Acoustic startle responses (ASR) at 6 dpf |  |  |  |  |  |  |  |  |  |  |  |  |  |  |  |  |  |  |
| --- | --- | --- | --- | --- | --- | --- | --- | --- | --- | --- | --- | --- | --- | --- | --- | --- | --- | --- |
| Parameter | ASR |  |  |  |  |  | SLC Kinematics |  |  |  |  |  |  |  |  |  |  |  |
|  | LLC Index |  | PPI |  | SLC Index |  | C1 Angle (degrees) |  | C1 Duration (s) |  | C2 Angle (degrees) |  | C2 Duration (s) |  | Distance (mm) |  | Latency (s) |  |
| Data | average | SD | average | SD | average | SD | average | SD | average | SD | average | SD | average | SD | average | SD | average | SD |
| 24- | 982.838 | 558.375 | 41.443 | 31.760 | 1311.261 | 540.625 | 130.123 | 12.015 | 10.632 | 0.967 | 83.123 | 15.467 | 10.053 | 1.248 | 77.000 | 8.917 | 7.930 | 1.425 |
| 24+ | **486.448** | **453.059** | 43.887 | 30.656 | 1203.887 | 580.210 | **100.784** | **32.508** | **10.059** | **1.720** | 83.827 | 19.030 | 10.431 | 2.427 | **57.962** | **14.113** | **10.212** | **2.133** |
| 48- | 1141.150 | 521.579 | 45.182 | 31.788 | 1255.768 | 422.756 | 136.774 | 10.670 | 11.581 | 0.872 | 87.484 | 16.335 | 11.226 | 1.679 | 71.323 | 8.494 | 8.290 | 1.197 |
| 48+ | **560.496** | **573.054** | 57.405 | 32.951 | 1325.044 | 717.360 | 129.500 | 26.400 | **11.115** | **1.121** | **108.577** | **22.430** | **13.308** | **1.659** | **56.731** | **8.786** | **7.192** | **2.602** |
| 72- | 1354.738 | 561.430 | 41.748 | 22.653 | 1370.996 | 447.377 | 142.167 | 10.319 | 11.708 | 0.789 | 90.750 | 16.604 | 11.792 | 2.020 | 72.208 | 9.147 | 8.042 | 1.457 |
| 72+ | 1152.673 | 609.306 | 49.760 | 29.732 | 1372.967 | 553.884 | 134.067 | 10.344 | **11.333** | **0.869** | 100.000 | 14.574 | 11.933 | 1.181 | 65.667 | 6.477 | 7.533 | 1.668 |

### Induction of dnVDR modulates early life stage acoustic startle response (ASR)

To further identify alterations in larval zebrafish sensory processing, we next assessed the acoustic startle response with six-day old zebrafish larvae that have been subjected to dnVDR induction as described with the VMR assay. Results of this assay (**Figure 2** and **Table 1B**) demonstrate a significant and specific impact on Long Latency C bends (LLC), quantified by calculating the area under the curve of stimulus intensity vs. LLC frequency (LLC index), post dnVDRa induction at both 24 and 48 hpf (p<0.0001 and p<0.001, respectively). Conversely, we observed that Pre-pulse Inhibition (PPI) and the total Short Latency C bends (SLC) responses, measured by the SLC index, were unaffected across all dnVDR induction time points. Kinematic assessment of SLC behaviors did reveal some significant alterations in the performance of these high-velocity escape responses at 24 and 48 hpf dnVDRa induction time points (**Figure 3**). For instance, the 24 hpf dnVDRa induced larvae exhibit a decrease in the initial C1 bend angle (p<0.0001), reduced C1 bend duration (p<0.05), increased overall latency (p<0.0001) and a decrease in distance moved (p<0.0001) for the total SLC response. The 48 hpf dnVDRa induced larvae exhibit an increased counterbend or C2 bend angle (p<0.0001) and increased C2 bend duration (p<0.0001), decreased overall latency (p<0.05) and a decrease in distance moved (p<0.0001) for the total SLC response as well. These results indicate that while dnVDR induced larvae fish do perform SLC responses at normal frequencies, their performance of that response is significantly impaired which may indicate that other deficits are present.

**Figure 2:**
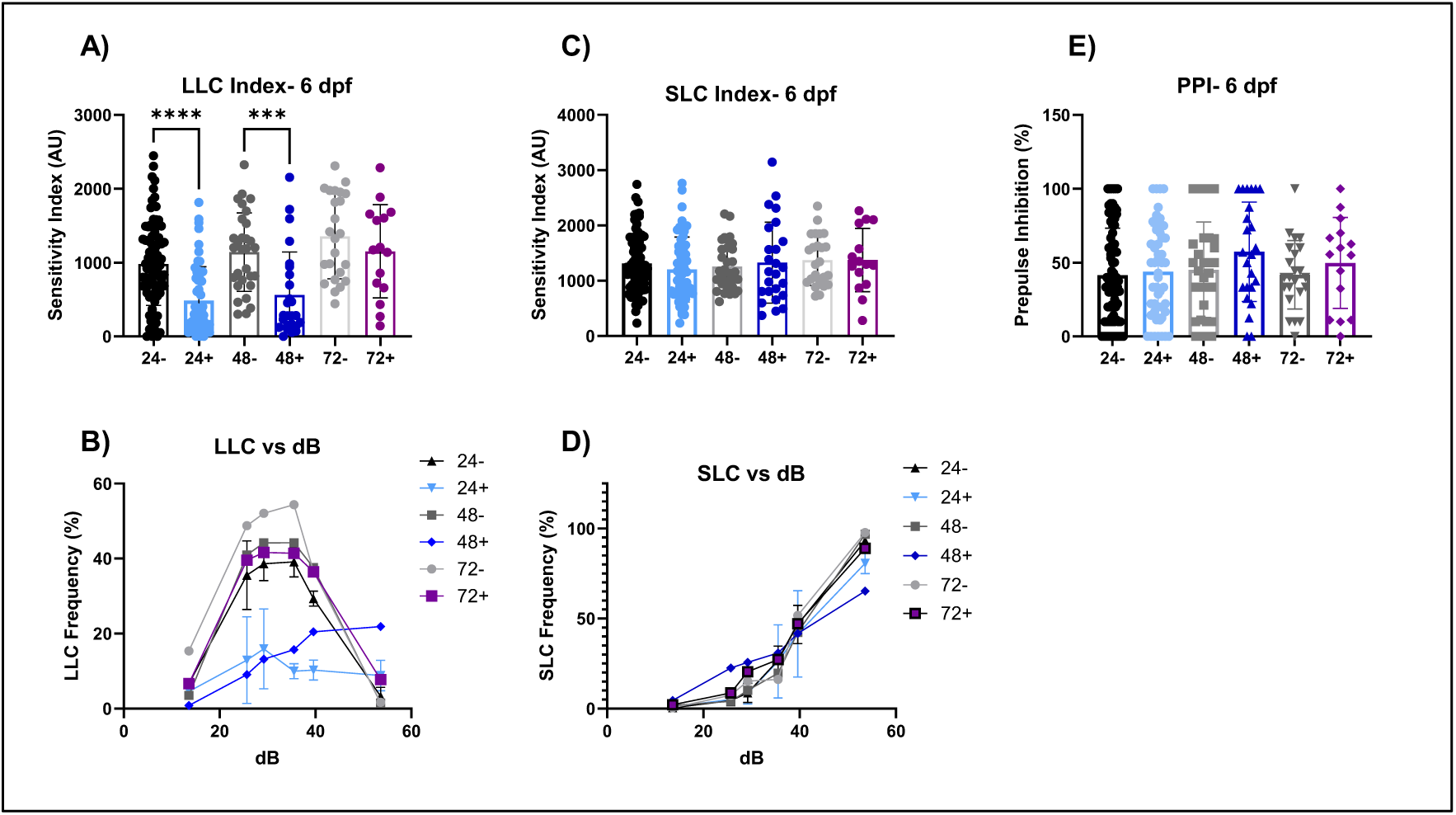
Acoustic Startle at 6 dpf. (A-B) These graphs represent the area under the curve of the LLC response frequency. There are significant decreases in LLC responses in the 24+ (p<0.0001) and 48+ (p≤ 0.001) fish. (C-D) These graphs represent the area under the curve of the SLC response frequency. There are no significant changes between groups for SLC response. (E) The percentage of PPI. We see no significant changes across the groups.

**Figure 3:**
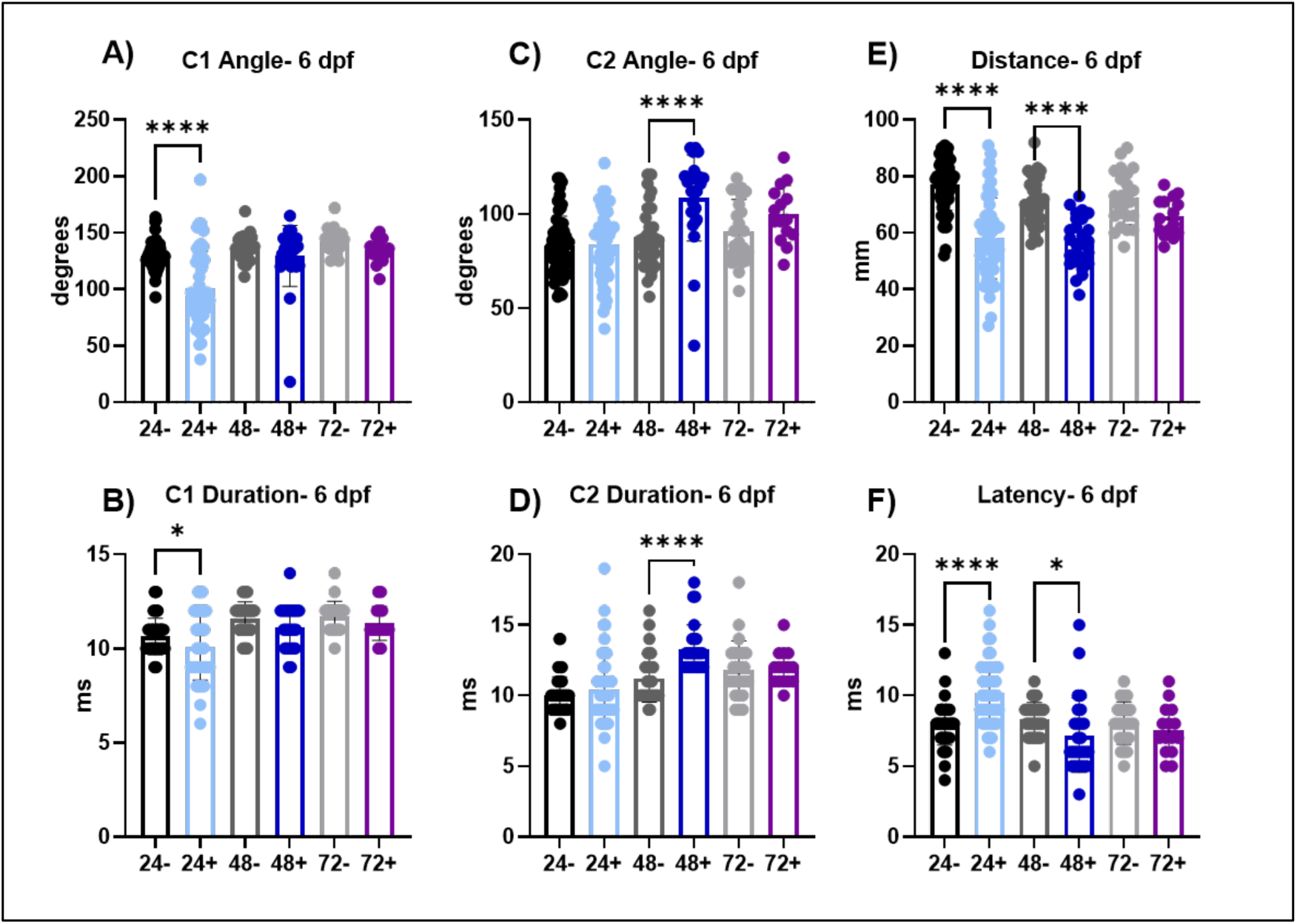
SLC response kinematics at 6 dpf. (A-B) C1 is the first C-bend performed. The angle and duration of the C1 bend is significantly decreased in 24+ fish (p<0.0001 and p<0.05) compared to the 24-fish. (C-D) C2 is the second C-bend performed. The angle and duration of C2 bend is significantly increased in 48+ fish (p<0.0001) compared to the 48-fish. (E-F) The total distance moved and latency, the total time of delay from the auditory stimulus, of the SLC response. The total distance of the SLC response is significantly decreased in 24+ fish and 48+ fish (p<0.0001) compared to their respective controls (24- and 48-). The latency is significantly increased in the 24+ fish (p<0.0001) compared to the 24-fish and significantly decreased in the 48+ fish (p<0.05) compared to the 48-fish.

### Induction of dnVDR results in persistent behavioral deficits in VMR and ASR assays

To assess if induction of dnVDRa expression during early development can result in long term deficits we assessed both behavior assays at 28 dpf (**Figures 4**-**6** and **Tables 2A-B**) following a single heat shock treatment between 24-72 hpf. With heat shock at 24 hpf, results indicate that deficits observed in the light dark locomotor activity assay at 6 dpf do not persist through 28 dpf. Conversely, when dnVDR is induced at 48 hpf, persistent effects on VMR outcomes are observed at the 28 dpf assessment. These include alterations in total distance moved (p<0.05 dark phase only), velocity (p<0.05 dark phase only), activity state (p<0.0001 and p<0.05, dark and light phases respectively), and distance to center point (p<0.05 and p<0.05, dark and light phases respectively). With induction of dnVDR occurring at 72 hpf and behavioral assessment conducted at 28dpf, we additionally observed persistent alterations in total distance moved (p<0.05), velocity (p<0.01), and activity state (p<0.05), however these observations occurred in the dark phase only.

**Figure 4:**
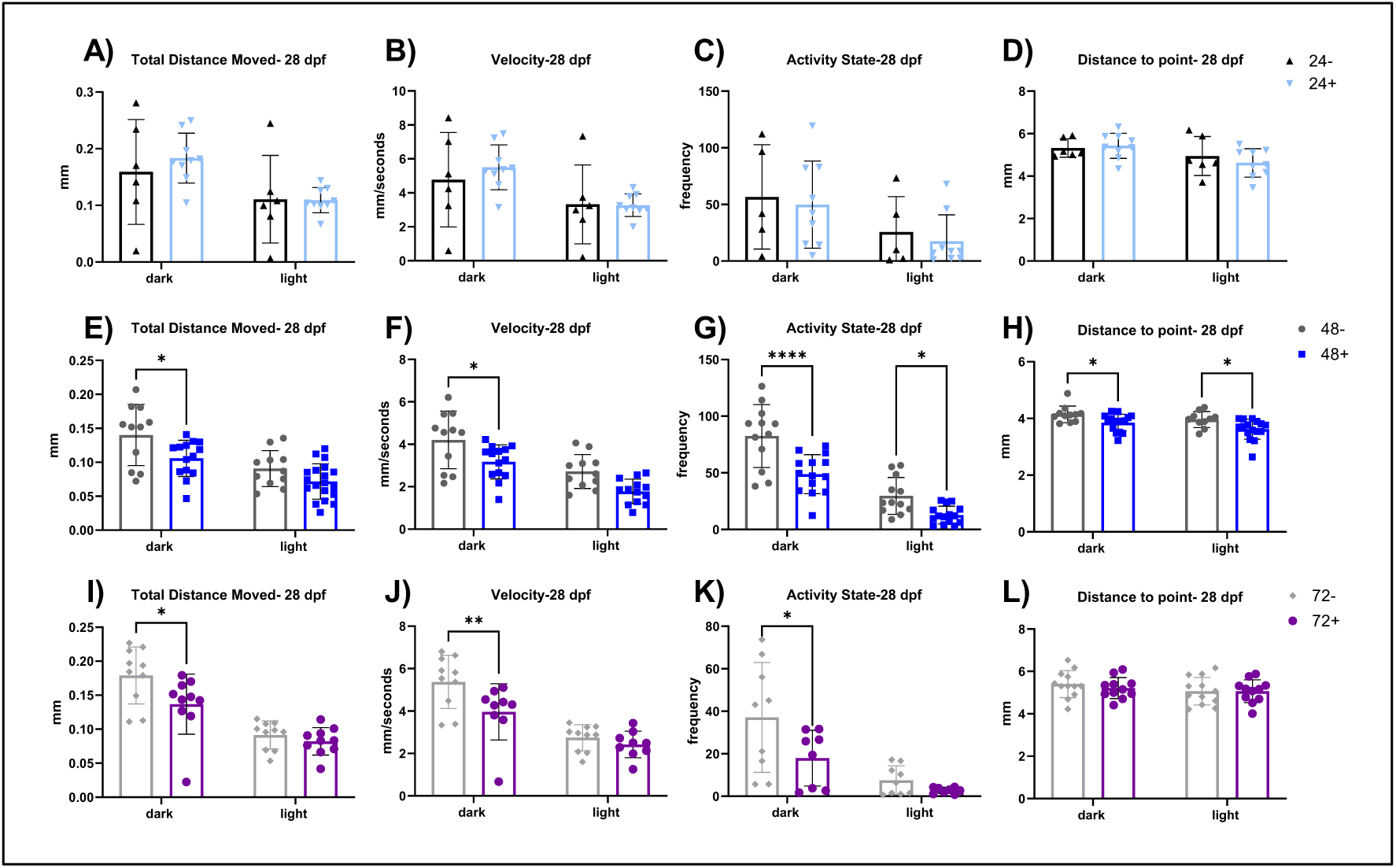
Visual motor response assay at 28 dpf. (A-D) VMR parameters in dnVDRa induced zebrafish at 24 hpf, there is no signficance difference at 28 dpf. (E-H) VMR parameters in dnVDRa induced zebrafish at 48 hpf. There is a significant decrease in distance moved in the dark period (p<0.05), a significant decrease in velocity in the dark period (p<0.05), a significant decrease in activity state in the dark period (p<0.0001) and light period (p<0.05) and a significant decrease in distance to point in the dark period (p<0.05) and light period (p<0.05) in the 48+ fish. (I-K) VMR parameters in dnVDRa induced zebrafish at 72 hpf. There is a significant decrease in distance moved in the dark period (p<0.05), a significant decrease in velocity in the light period (p<0.01) and a significant decrease in activity state the light period (p<0.05) in the 72+ fish.

**Table 2:** All behavioral data values at 28 dpf. (2A) Data values for all dnVDRa induced timepoints (24+, 48+ and 72+) and their controls (24-, 48-, 72-) for all VMR parameters. (2B) Data values for all dnVDRa induced timepoints (24+, 48+ and 72+) and their controls (24-, 48-, 72-) for all ASR parameters. Bolded values represent statistical difference between the dnVDRa induced fish versus their controls.

| Table 2A: Visuo-motor response (VMR) assay at 28 dpf |  |  |  |  |  |  |  |  |  |  |  |  |  |  |  |  |
| --- | --- | --- | --- | --- | --- | --- | --- | --- | --- | --- | --- | --- | --- | --- | --- | --- |
| Parameter | Distance Moved (mm) |  |  |  | Velocity (mm/s) |  |  |  | Activity State (frequency) |  |  |  | Distance Moved (mm) |  |  |  |
| Bin | Dark |  | Light |  | Dark |  | Light |  | Dark |  | Light |  | Dark |  | Light |  |
| Data | average | SD | average (mm) | SD | average (mm/s) | SD | average (mm/s) | SD | average (freq) | SD | average (freq) | SD | average (mm) | SD | average (mm) | SD |
| 24- | 0.159 | 0.085 | 0.111 | 0.071 | 4.772 | 2.538 | 3.320 | 2.120 | 56.590 | 41.196 | 25.592 | 27.975 | 5.324 | 0.394 | 4.947 | 0.836 |
| 24+ | 0.183 | 0.042 | 0.109 | 0.021 | 5.500 | 1.246 | 3.273 | 0.630 | 49.867 | 36.295 | 17.451 | 22.034 | 5.428 | 0.557 | 4.622 | 0.632 |
| 48- | 0.140 | 0.043 | 0.091 | 0.025 | 4.205 | 1.292 | 2.719 | 0.762 | 82.506 | 26.680 | 29.635 | 15.617 | 4.145 | 0.280 | 3.962 | 0.268 |
| 48+ | 0.106 | 0.026 | 0.072 | 0.025 | 3.171 | 0.771 | 1.777 | 0.554 | 48.830 | 16.625 | 12.736 | 7.513 | 3.850 | 0.287 | 3.620 | 0.343 |
| 72- | 0.179 | 0.040 | 0.091 | 0.019 | 5.372 | 1.193 | 2.741 | 0.584 | 37.097 | 24.361 | 7.482 | 6.400 | 5.400 | 0.607 | 5.066 | 0.623 |
| 72+ | 0.137 | 0.042 | 0.082 | 0.019 | 3.960 | 1.248 | 2.426 | 0.591 | 17.884 | 12.293 | 2.725 | 1.336 | 5.213 | 0.476 | 5.065 | 0.516 |

**Table 2:** All behavioral data values at 28 dpf.
| Table 2B: Acoustic startle responses (ASR) at 28 dpf |  |  |  |  |  |  |  |  |  |  |  |  |  |  |  |  |  |  |
| --- | --- | --- | --- | --- | --- | --- | --- | --- | --- | --- | --- | --- | --- | --- | --- | --- | --- | --- |
|  | ASR |  |  |  |  |  | SLC Kinematics |  |  |  |  |  |  |  |  |  |  |  |
| Parameter | LLC Index |  | PPI |  | SLC Index |  | C1 Angle (degrees) |  | C1 Duration (s) |  | C2 Angle (degrees) |  | C2 Duration (s) |  | Distance (mm) |  | Latency (s) |  |
| Data | average | SD | average | SD | average | SD | average | SD | average | SD | average | SD | average | SD | average | SD | average | SD |
| 24- | 332.625 | 160.679 | 56.974 | 32.588 | 415.531 | 136.122 | 130.733 | 16.611 | 11.688 | 2.083 | 88.375 | 24.990 | 15.933 | 6.688 | 164.688 | 19.852 | 9.067 | 1.652 |
| 24+ | 286.538 | 144.083 | 62.468 | 32.543 | 552.310 | 170.710 | 121.045 | 17.134 | 11.304 | 1.755 | 90.818 | 23.346 | 12.955 | 4.311 | 190.864 | 37.165 | 6.696 | 1.572 |
| 48- | 325.817 | 140.258 | 81.527 | 21.383 | 524.594 | 101.110 | 137.429 | 33.746 | 11.190 | 4.238 | 77.714 | 31.545 | 10.429 | 3.762 | 174.048 | 21.973 | 7.789 | 2.440 |
| 48+ | 266.353 | 157.493 | 47.498 | 25.930 | 602.796 | 153.942 | 142.333 | 29.323 | 10.591 | 1.527 | 89.200 | 15.237 | 10.190 | 1.842 | 190.800 | 24.447 | 6.250 | 1.841 |
| 72- | 303.146 | 156.440 | 92.739 | 9.681 | 532.959 | 133.344 | 125.000 | 21.312 | 10.818 | 0.886 | 77.238 | 18.231 | 10.095 | 1.540 | 176.667 | 14.936 | 6.565 | 2.039 |
| 72+ | 241.312 | 165.142 | 58.011 | 26.449 | 649.491 | 189.610 | 125.750 | 12.365 | 10.500 | 0.806 | 84.474 | 16.678 | 10.579 | 2.085 | 180.000 | 23.906 | 6.650 | 1.590 |

We did not observe significant alterations in LLC index at 28 dpf (**Figure 5A**) with dnVDRa induced in zebrafish embryos at any early developmental stage (24, 48 or 72). We did however observe an increase in SLC index in the 24 hpf dnVDRa induced fish (p<0.05) and the 72 hpf dnVDRa induced fish (p<0.05). We did not observe persistent alterations in the C-bends when we analyzed SLC kinematics at 28 dpf (**Figure 6**). We did however observe an increase in distance moved in the 24 hpf dnVDRa induced fish (p<0.01) and a decrease in latency in both 24 and 72 hpf dnVDRa induced fish (p<0.001 and p<0.05, 24+ and 48+ respectively) in the SLC response. We also observed a significant decrease in pre-pulse inhibition (PPI) in our 48 hpf dnVDRa induced fish (p<0.0001) and 72 hpf dnVDRa induced fish (p<0.001) when assayed at 28 dpf (Figure 5C). Pre-pulse inhibition (PPI) is a key form of sensorimotor plasticity in which a weak pre-pulse reduces responses to subsequent strong stimuli. Interestingly PPI was unaffected when early-stage embryos were induced and assayed at 6dpf (**Figure 2E**). However, as with the VMR assay, 28-day-old young adult fish that received a single heat shock at 48 hpf displayed significantly reduced PPI, indicating dysfunction in sensory processing later in development.

**Figure 5:**
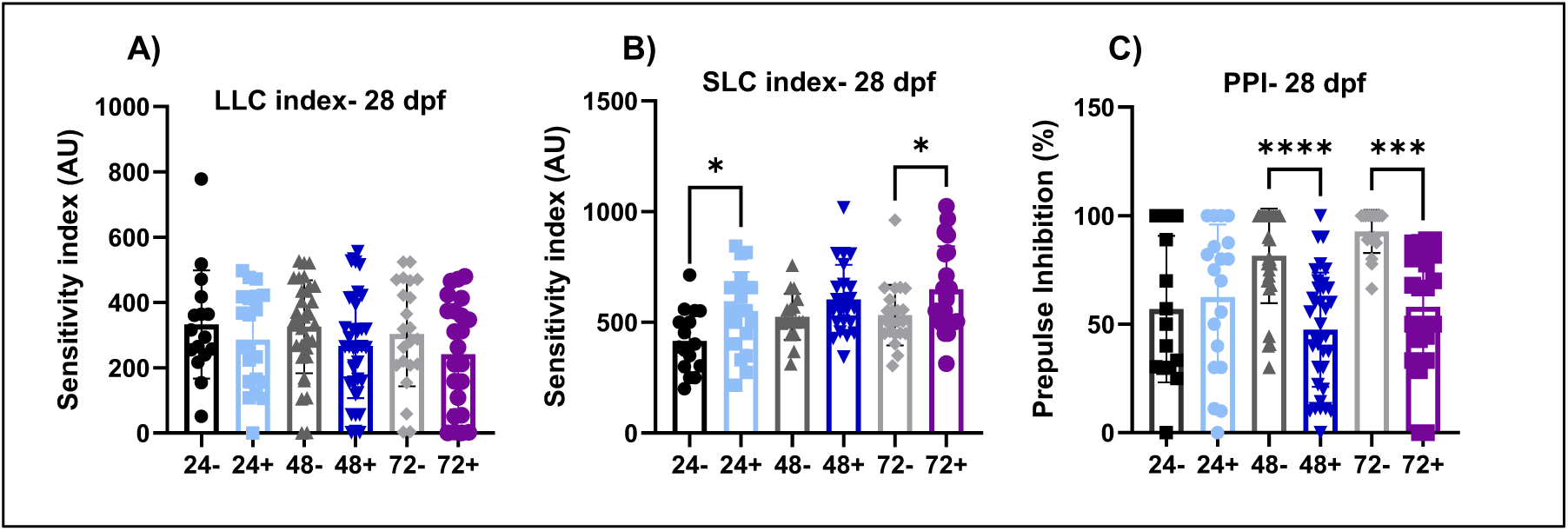
Acoustic Startle at 28 dpf. (A-B) These graphs represent the area under the curve of the SLC response frequency and LLC response frequency, respectively. There is a significant increase in SLC responses in the 24+ (p≤ 0.05) and 72+ (p≤ 0.05) fish. (C) We can see a significant decrease of PPI in the 48+ and 72+ fish (p<0.0001).

**Figure 6:**
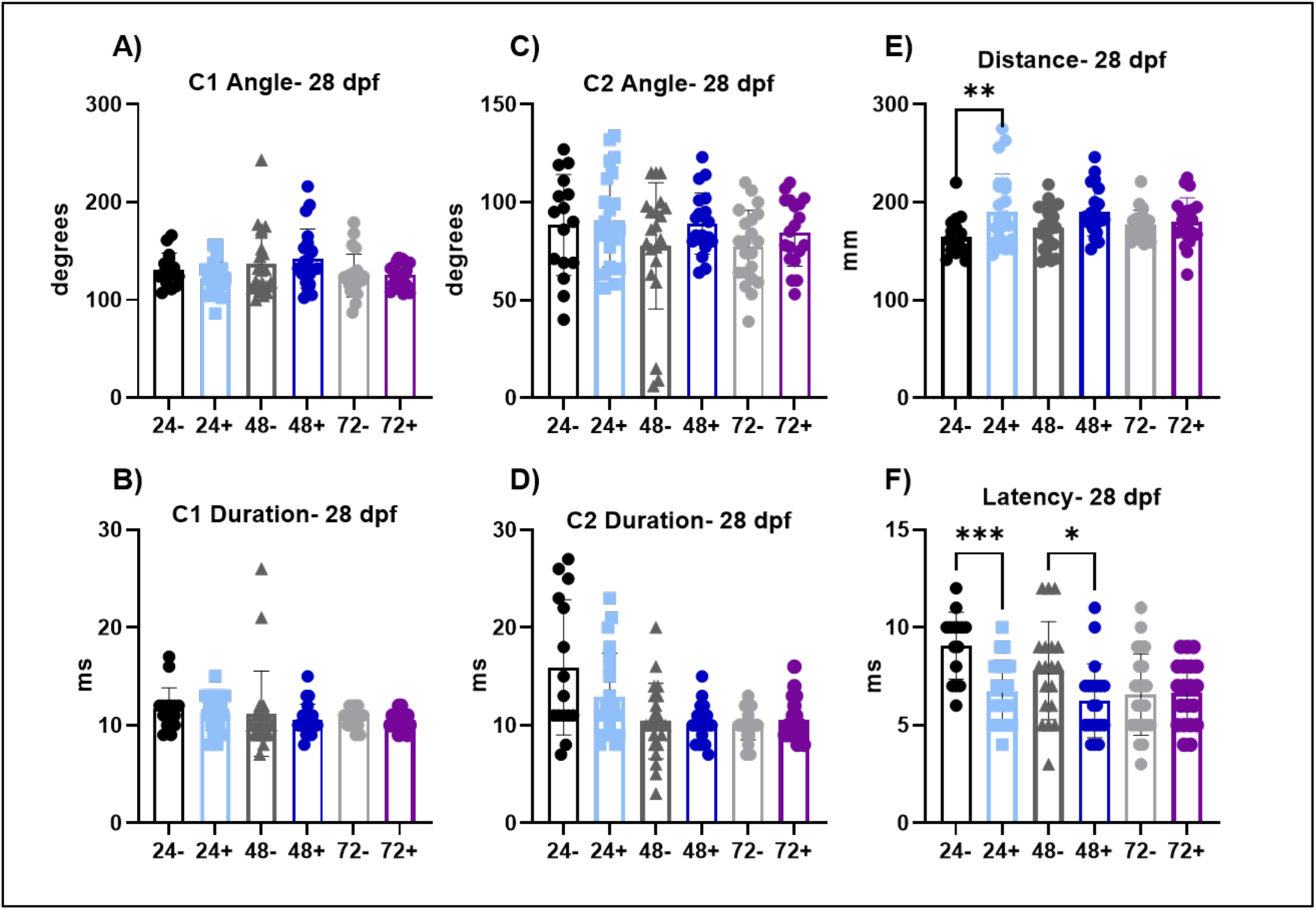
SLC response kinematics at 28 dpf. (A-B) C1 is the first C-bend performed. The angle and duration of the C1 bend is not significantly altered at this time point. (C-D) C2 is the second C-bend performed. The angle and duration of C2 bend is not significantly altered at this time point. (E-F) The total distance moved and latency, the total time of delay from the auditory stimulus, of the SLC response. The total distance of the SLC response is significantly increased in 24+ fish (p<0.01) compared to 24-fish. The latency is significantly decreased in the 24+ fish (p<0.001) and the 48+ fish (p<0.05) compared to their respective controls (24- and 48-).

### Neurotransmitter gene expression at 6 dpf

Next, we examined expression of genes associated with six select neurotransmitter pathways with known roles in sensory function; cholinergic, glycinergic, serotonergic, GABAergic, glutaminergic and dopaminergic. Results from this assessment indicate significant alterations in expression of multiple genes across each dnVDRa induction period and within all pathways examined (**S Figure 2**). In order to visualize global gene expression events within these pathways, gene expression was formatted into a heat map using hierarchical clustering (**Figure 7**). Gene expression data resulted in two empirical clusters, with C-I comprised of the 48hpf dnVDRa induction period, and C-II comprised of two sub clusters including the 24hpf and the 72hpf induction period. Results indicate that gene expression with the 48 hpf dnVDRa induced fish was significantly different from the 24 and the 72 hpf dnVDRa induced fish, which are more similar to the control group.

**Figure 7.**
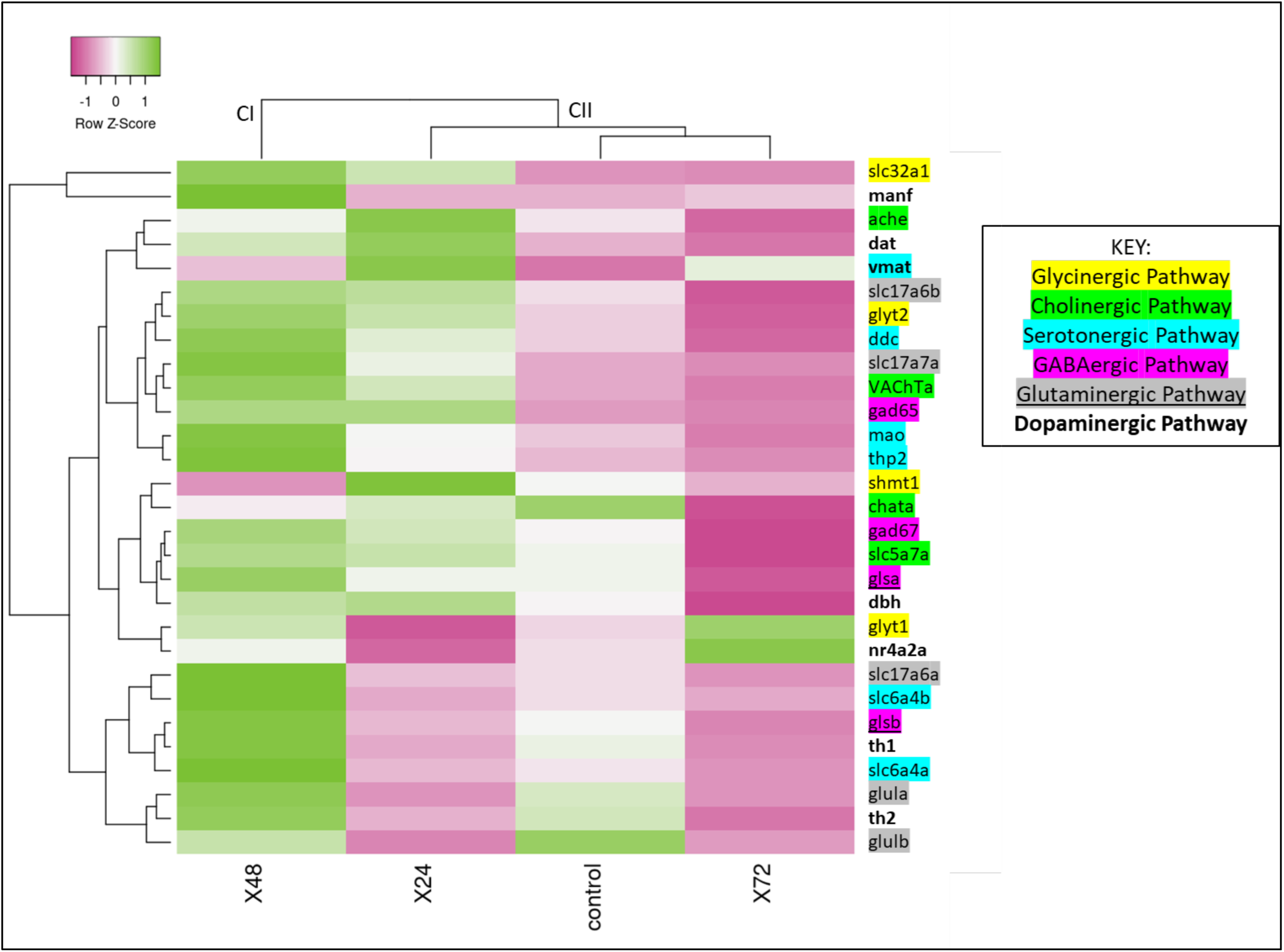
Consolidated gene expression of neurotransmitter pathways. Heat map showing the row Z-score, a scaling method to compare all values in the row, for gene expression in neurotransmitter genes at 6 dpf in dnVDRa induced fish at either 24, 48 or 72 hpf and the control group. Higher gene expression values are indicated in green while lower values are in pink. The key depicts what genes belong to what pathway. Genes that fall into more than one pathway are indicated by color and either underline or bold.

## Discussion

The nervous system is able to distinguish between different sensory stimuli, integrate multiple simultaneous stimuli and processes the appropriate behavioral response in specific contexts. Specifically, in vertebrates visual, tactile, and auditory stimuli all elicit stereotyped behaviors[40–43]. These behavioral responses are modulated by specific parameters associated with each stimuli including luminance, size, and speed (visual)[44–46] or intensity (tactile, auditory)[43, 47]. The importance of proper sensory processing and proper behavioral responses can be highlighted by the findings that disruptions in sensory responsiveness are observed in many neuropsychiatric disorders including anxiety[48–51], PTSD[52], ADHD[53], migraine[54], schizophrenia[55, 56], and autism[57–59]. The molecular and cellular mechanisms that drive the development of the neural circuits regulating sensorimotor responsiveness are complex and incompletely understood. In this study we observe that modulation of VDR signaling during early zebrafish neurodevelopment results in attenuation of sensory processing activities. Specifically, we observe alterations in defined locomotion activities within our light/dark (VMR) assay and the ability to perform and execute Long Latency C-bend (LLC) response and Short Latency C-bend (SLC) responses in our acoustic startle response assay. These studies demonstrate that early embryonic disruptions in vitamin D signaling modulate sensory processing activities with lasting effects in juvenal forms consistent with a Developmental Origins of Health and Disease (DOHaD)[26, 60] model of neurodevelopment.

The VMR assay is a well-established test that measures swimming behaviors in response to whole-field light/dark stimuli[61, 62]. For zebrafish larvae, transition from light to dark results in increased locomotion (velocity, distance moved, mobility, activity state, and angular velocity), typically attributed to a stressed- or anxiety-like state, while transition from dark to light results in decreased locomotion. This behavioral plasticity requires proper functioning of visual and motor systems, as well as integration centers in the brain that connect visual perception, internal state, and locomotor function[63]. Overall, our assessments suggest that modulation of VDR signaling through induction of a dominant form of VDR (dnVDR) at 24 or 48 hpf resulted in marked decrease in both activity state and hypolocomotion in both light and dark phases of our locomotor activity assay at 6 dpf. When we assess this behavior at 28 dpf, we demonstrate that dnVDRa induction at 48 or 72 hpf results in a significant decrease in all the tested locomotor parameters. Induction of the dnVDRa at 48 hpf caused the most severe and persistent behavioral phenotypes, eliciting sustained hypolocomotion in both light and dark phases at 28 dpf. This reduced locomotion in the light-dark transitions suggest that disrupting developmental VDR signaling leads to impaired sensorimotor integration. Interestingly, contrary to our findings, studies in rodent models have demonstrated hyperlocomotion with developmental vitamin D deficiency[64]. Our model, however, utilizes a conditional dominant negative VDR knock down system that enables temporal regulation of VDR and behavior assessment at very early stages of organismal development compared to rodent models of VDD. Overall, our findings confirm previous observations from our group that dietary modulations or environmental exposures to xenobiotics that alter developmental VDR signaling results in altered behavioral function[65, 66].

We next tested zebrafish startle responses using an acoustic startle response assay. In this assay zebrafish are exposed to an abrupt acoustic stimulus and can respond with one of two stereotyped behaviors either Short-Latency C-bends (SLCs) or Long-Latency C-bends (LLCs). These responses have been identified by multiple kinematic parameters such as latency, turn angle, and pectoral fin involvement. SLCs are typically performed when zebrafish are exposed to intense acoustic stimuli and are initiated within 4-15 ms. Weaker acoustic stimuli typically trigger LLCs and are initiated 15-100 ms after the stimulus, are more varied in terms of kinematics, and can be influenced by visual input[67, 68]. Additionally, these behaviors have been demonstrated to have differing neural circuitry. The SLC response is triggered by Mauthner neurons (M-cells), which receive direct input from auditory afferents and contact spinal motor neurons[38, 68, 69]. LLC responses occur independent of M-cell function and instead depend on a bilateral population of pre-pontine neurons located between the locus coeruleus and cerebellum[68]. It has been hypothesized that due to the delayed response, LLC behaviors may allow for integration of multimodal stimuli and context-appropriate decision-making[68].

In this study we find that early developmental induction of dnVDR between 24-48 hpf in zebrafish embryos causes a severe reduction in LLC responses but relatively little effect on SLC responses. These data suggest that VDR signaling selectively impacts LLC and not SLC circuit function, as all groups are able to perform SLC reactions at normal rates. However, our data also indicate that dnVDRa induction at 24 or 48 hpf alters SLC kinematics at 6 dpf. Specifically, dnVDRa induction at 24 hpf reduces C1 bend angle and duration while dnVDRa induction at 48 hpf increases C2 bend angle and duration. Induction at both 24 and 48 hpf reduced the distance traveled during SLC responses at 6 dpf, and so together these findings indicate that while VDR signaling has minimal impacts on SLC initiation, it is important for the fine motor control of these responses, likely affecting spinal motor circuits. Conversely, the long-term effects of developmental dnVDR induction on the acoustic startle response in 28 dpf fish were mostly reversed. LLCs were unaffected while SLCs were increased in the 24 hpf and 72 hpf groups. We also observed pronounced deficits in prepulse inhibition when dnVDRa is induced at 48 or 72 hpf. Pre-pulse inhibition (PPI) is a neurological phenomenon in which a weak pre-pulse reduces responses to subsequent strong stimuli and can be related to sensorimotor plasticity. PPI can also be a measure for sensorimotor gating, which can assess the brain’s ability to filter out irrelevant sensory information and prevent an excessive response to stimuli. PPI has become important in the context of neuropsychiatric disorders and could be a potential biomarker of brain function[69, 70]. Deficits in PPI are a common endophenotype of schizophrenia[55, 56], so this finding is consistent with the association of VDD with schizophrenia[71, 72] and further supports the importance of investigating the underlying mechanisms. The deficits in PPI at 28 dpf may indicate an inability to properly modulate startle responses later in development. With these behavioral assays we demonstrate that dnVDRa induction at 48 hpf results in the most severe behavior phenotypes that are sustained to 28 dpf, highlighting the importance of VDR signaling during development for both visually-driven locomotor activity and acoustically-evoked behaviors and behavioral modulation. Together our behavior data reveal that developmental disruption of VDR signaling causes multiple yet specific defects in neural function (e.g. LLCs but not SLCs affected). This suggests that the defects are unlikely to be secondary to systemic effects of VDR dysfunction but are rather the result of the loss/modulation of VDR function in the underlying neural circuits. This is further reinforced by the fact that the dnVDR is largely expressed in the brain and spinal cord, with comparatively little expression in non-neural tissues in our zebrafish model(S Figure 1).

To coordinate behavior phenotypes with putative molecular changes caused by dnVDR induction, we examined gene expression in multiple neurotransmitter pathways in the context of development. In addition to driving all sensory and motor function, neurotransmitter signaling impacts cell proliferation, migration, and differentiation during brain development[73–79]. Vitamin D is also well-known to modulate synaptic function by altering the expression of receptors, transporters and the enzymes involved in metabolism and synthesis of neurotransmitters[13, 64, 80–82]. Previous DVD rat models have demonstrated disruptions to dopaminergic, glutamatergic and serotonergic pathways[21, 64, 80, 81, 83–86]. Specifically, it has been demonstrated that VDR activation can result in induction in expression of tryptophan hydroxylase 2 (THP2) which is the rate limiting enzyme in the synthesis of serotonin[80] and has been found to repress expression of monoamine oxidase-A (MAO) and serotonin reuptake transporter (SERT) at certain concentrations *in vitro*[84]. These trends indicate vitamin D modulate the serotonergic system to result in increased serotonin levels. In the dopaminergic system, VDR overexpression resulted in elevated catechol-o-methyl transferase (COMT), the crucial enzyme in dopamine synthesis and *in vivo* vitamin D-deficient rats had reductions in COMT [21] indicating VDR signaling is important for dopamine synthesis. Within the inhibitory systems (GABA, glutaminergic), dietary vitamin D deficient rodent models identified that low vitamin D levels resulted in decreased expression of glutamate and GABA transporters (GAT)[85]. Additionally, in an adult vitamin D deficient mouse model, expression of glutamic acid decarboxylase (GAD65/67) an important enzyme in GABA synthesis was found to be reduced[81]. These studies indicate that VDR signaling is involved in both excitatory and inhibitory pathways in the brain. In this study we focused on key components of cholinergic, GABAergic, glycinergic, glutamatergic, serotonergic, and dopaminergic neurotransmitter pathways at 6 dpf in all dnVDR induced groups compared to our non-induced dnVDR controls. Our data demonstrate that induction of dnVDR modulates expression of essential genes within these pathways. Expression differed dependent upon the timing of dnVDRa induction (**Figure 8** and **S Figure 2**). Consistent with our behavior data, dnVDRa induction at 48 hpf resulted in gene expression changes that were most different from control groups and the other induction time points when analyzed through hierarchical clustering. These data are consistent with the current findings in mammalian models of VDD and illustrate the importance of temporal disruptions in vitamin D signaling and modulation of essential neurotransmitter pathways. These broad impacts likely contribute to the persistent behavior phenotypes observed in this study.

These studies provide insight on how developmental VDR signaling modulates neural gene expression and behavior. In conjunction, mammalian model systems of DVD depict distinct behavioral phenotypes such as significant alterations in social behavior, learning and memory. As such, current research supports the hypothesis that developmental vitamin D deficiency adversely impacts brain development resulting in modulation of adult behaviors consistent with a DOHaD model of VDD[26, 60]. Yet a significant gap remains in our understanding of both the mechanisms of VDD on DOHaD and windows of susceptibility to VDD.

## Supporting information

Ritter et al 2025

## Acknowledgements

The authors would like to acknowledge the following funding sources for support on this project: NIEHS T32ES07046 Molecular Pathways to Pathogenesis in Toxicology, NIEHS P30ES025128 Center for Human Health and the Environment, P42 ES031009-01 National Institute for Environmental Health Sciences Center for Health and Environmental Effects of PFAS, and NINDS R01NS116354.

