## Supplementary material for "Early disruptions in vitamin D receptor signaling induces persistent developmental behavior deficits in zebrafish larvae": Ritter et al 2025

**Supplemental Table 1.** Primer sequences for neurotransmitter related qPCR.

| Gene | Primer Sequence (5'- 3') | Product Size (bp) |
|---|---|---|
| ache | TCCTGCTTTTGCTTTTCGCC (forward) | 245 |
| | AACGTGAGTAGCACAGTGGG (reverse) | |
| chata | ACGAGTCGACAACATTCGCT (forward) | 114 |
| | ATGGCAGTCCAGAGCAACTC (reverse) | |
| dat | CTAAAAAGCTCCGCATCCAG (forward) | 231 |
| | TGTCCAAGAGCAAAGCAATG (reverse) | |
| dbh | TGGCCTATCATATCCCGCTG (forward) | 209 |
| | CCACGCATCCCCAAAATAGG (reverse) | |
| ddc | ACTGGCACAGCCCATACTTC (forward) | 169 |
| | AGCATCTTTCCCAGCCAGTC (reverse) | |
| ef1a | TACAAATGCGGTGGAATCGAC (forward) | 246 |
| | GTCAGCCTGAGAAGTACCAGT (reverse) | |
| gad65 | ATGGTGCCTTTGATCCTTTG (forward) | 228 |
| | TCTGCATCAGTCCCTCCTCT (reverse) | |
| gad67 | GACGACAAGGGTCGAATTGT (forward) | 211 |
| | TGCGCACGTAGTTAGTGAGG (reverse) | |
| glsa | TTTAGGCTTTCGAGGGCGTT (forward) | 165 |
| | GCCTGCCTTCCTTCTCTTGT (reverse) | |
| glsb | TGTGGACTAAAGTGTCGCCC (forward) | 175 |
| | ACGTGACCCGAATGAGAACC (reverse) | |
| glula | TGCCTCAGGGAGACCAAGTA (forward) | 158 |
| | GCCTTCAGCTTGATACGTGC (reverse) | |
| glulb | ATATCAGGCTGAGGGGTCCA (forward) | 272 |
| | GAGGTCCAGGGAAGCCATTG (reverse) | |
| glyt1 | TGAGCACTCATCGTGCCAAT (forward) | 242 |
| | GCCTGATGACTTGACTCCCC (reverse) | |
| glyt2 | TTCCAGGACGATGATGATGA (forward) | 237 |
| | GAACTGACCCAACGACACCT (reverse) | |
| manf | AGCGGCTTTACCAGACGATA (forward) | 149 |
| | CTTTGTTGCTGCATCACTCG (reverse) | |
| mao | TCTGGCACGAAAATCACGGA (forward) | 103 |
| | TAACGCCTCCTCTGATCCCA (reverse) | |

**Supplemental Table 1**. Primer sequences for neurotransmitter related qPCR. (continued)

| | | |
|---|---|---|
| nr4a2a | AGTATGGCTCATCTCCGCAA (forward) | 209 |
| | CGGCTTCACGTCATAACTGG (reverse) | |
| shmt1 | CTTCTGGCAGACATGGCTCA (forward) | 280 |
| | GCTTCAAAGCAACAGCGACA (reverse) | |
| slc17a6a | GCCACTCTGCTGTTAGTGGT (forward) | 184 |
| | TACCAGAAAGTGTGCCGACC (reverse) | |
| slc17a6b | TCCATGCCCGTCTATGCAATC (forward) | 205 |
| | TTTTGCTGCGAAGGTGATCG (reverse) | |
| slc17a7a | CGGCTCATTCTTCTGGGGTT (forward) | 166 |
| | GACCATGATCACACACCCGT (reverse) | |
| slc32a1 | CGGACAAGCCCAGAATCACT (forward) | 228 |
| | CGTACGAGTCTCTCACTCGC (reverse) | |
| slc5a7a | GAAGCTGGTGGACTCCTGAT (forward) | 262 |
| | TGTCCCCTGTGAGATAAAAATTCCC (reverse) | |
| slc6a4a | CTCCAGGCTCTCTATCCCCT (forward) | 188 |
| | CGGAACGCTATCCACAACCA (reverse) | |
| slc6a4b | AGCACTTTTGGTGGGTTGGA (forward) | 229 |
| | AGGAGACTGCTGTTGCTTCC (reverse) | |
| th1 | AGTGTGAAGTGCACCTGTCG (forward) | 233 |
| | GGCAATGTCTCCGATCATCT (reverse) | |
| th2 | CCCAGTACATTCGTCACCCT (forward) | 238 |
| | GCCCCATAAGCCTTTACTGC (reverse) | |
| thp2 | AGCCCCTAAATCACCTGTTGG (forward) | 180 |
| | TTAAAGGATGTCCTGCGGAGC (reverse) | |
| VAChT | GCCGAGGCTTCACTATTCAG (forward) | 223 |
| | GTCCTGGATCGCATTTCCTA (reverse) | |
| vmat | TCTTTCTCCATCGAGCACCT (forward) | 202 |
| | GGCAAACAACAGACCCACTT (reverse) | |

**Supplemental Table 1**. Primer sequences for neurotransmitter related qPCR. Gene expression was normalized to *efla*, the housekeeping gene.

**Supplemental Figures**

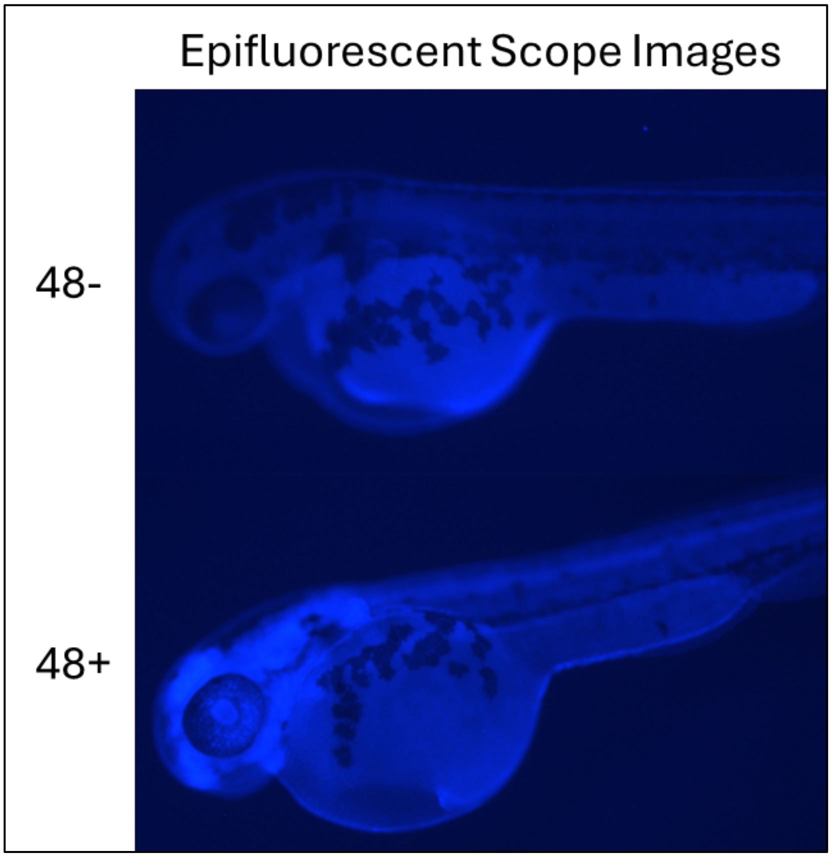


**S Figure 1.**BFP expression in 48 hpf heat shocked transgenic zebrafish. Epifluorescent images of TG(hsp:dnVDRa:BFP) heat shocked at 48 hpf and imaged 5 hours post heat shock. We can see fish heat shocked but very minimal BFP expression which are deemed 48-. The dnVDRa positive fish, 48+, have distinct BFP expression in the cranial region.

SF1A-Glycinergic Pathway Genes

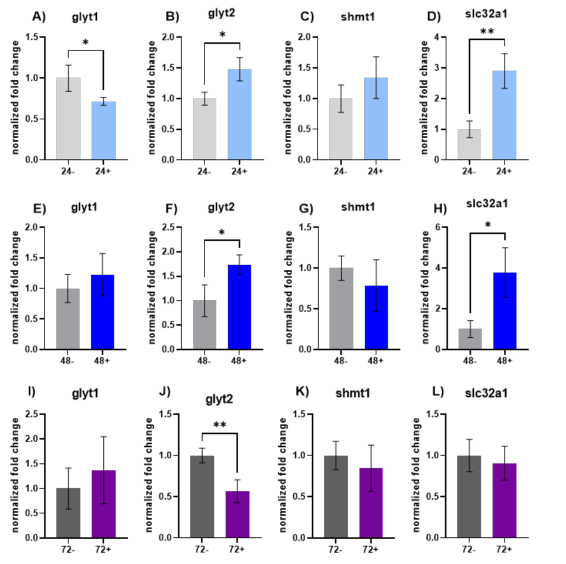


SF1B-Cholinergic Pathway Genes

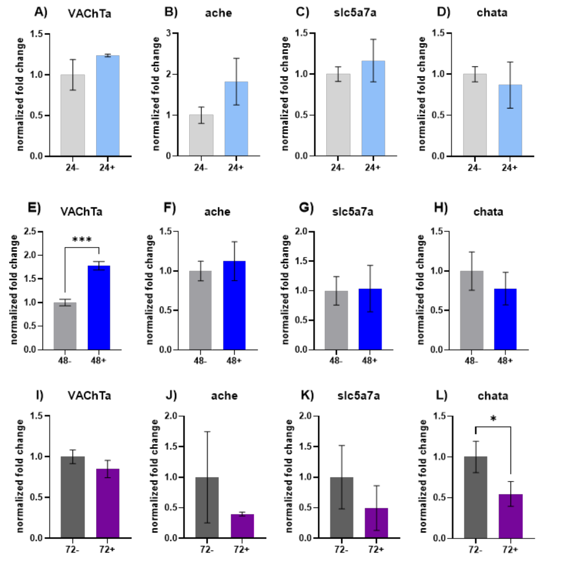


SF1C-Serotonergic Pathway Genes

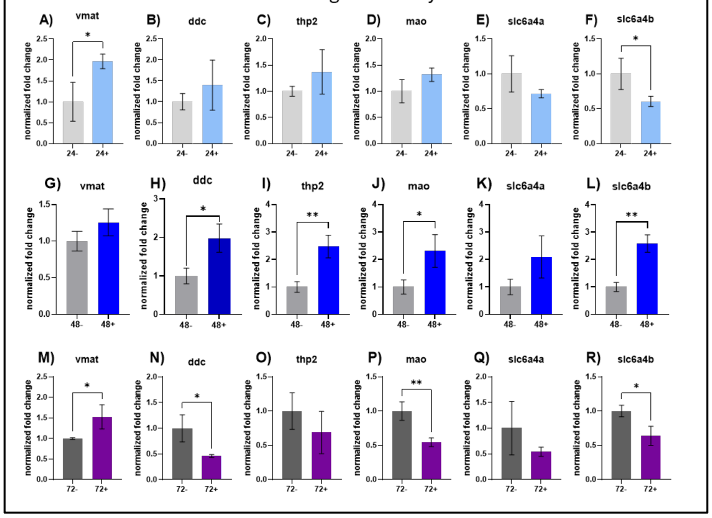


SF1D-GABAergic Pathway Genes

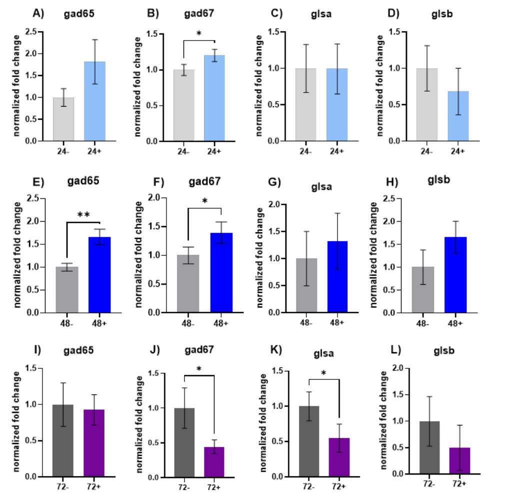

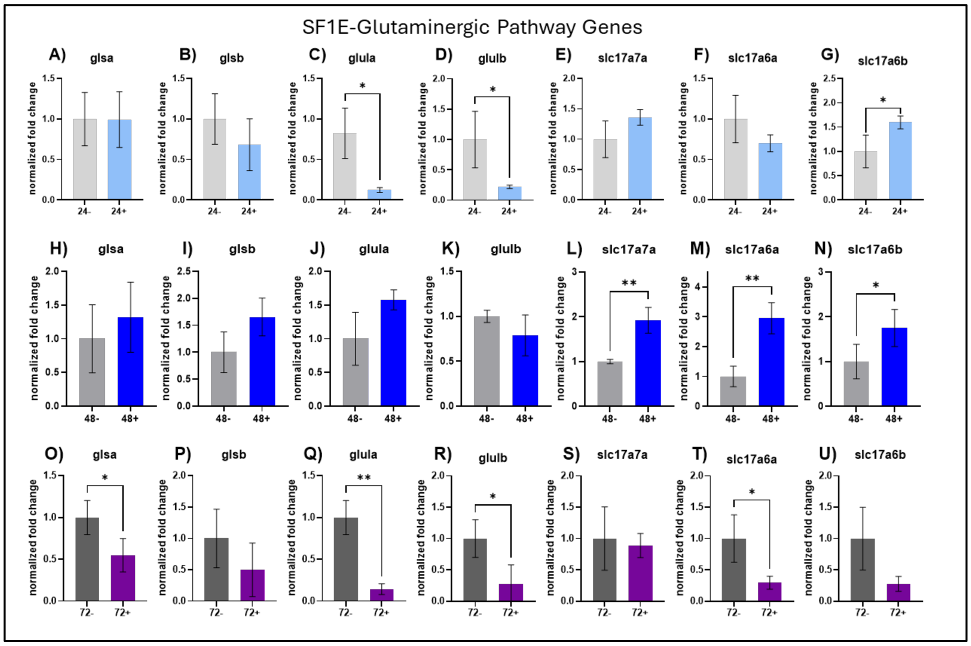


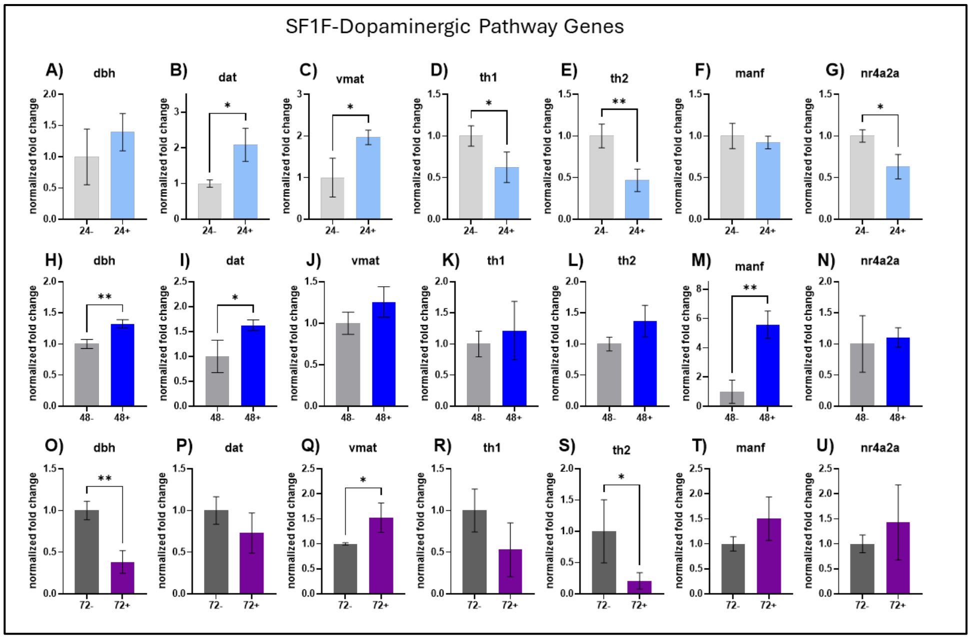


**Supplementary Figure 2. Gene expression of neurotransmitter pathways at 6 dpf. SF1A)** Glycinergic pathway gene expression. In the 24 hpf dnVDRa induced group we see a significant decrease in glyt1 expression ($p<0.05$) and an increase in glyt2 expression ($p<0.05$) and in slc32a1 expression ($p<0.01$). In the 48 hpf dnVDRa induced fish there is a significant increase in glyt2 and slc32a1 expression ($p<0.05$). In the 72 hpf dnVDRa induced fish there is a significant decrease in glyt2 expression ($p<0.01$). **SF1B)** Cholinergic pathway gene expression. There are no altered genes in this panel for the 24 hpf dnVDRa induced fish. In the 48 hpf dnVDRa induced fish there is a significant increase in VAChTa expression ($p<0.001$). In the 72 hpf dnVDRa induced fish there is a significant decrease in glyt2 expression ($p<0.05$). **SF1C)** Serotonergic pathway gene expression. In the 24 hpf dnVDRa induced group we see a significant decrease in slc6a4b expression ($p<0.05$). In the 48 hpf dnVDRa induced fish there is a significant increase in ddc ($p<0.05$), thp2 ($p<0.01$), mao ($p<0.05$) and slc6a4b ($p<0.01$) expression. In the 72 hpf dnVDRa induced fish there is a significant decrease in ddc ($p<0.05$), mao ($p<0.01$) and slc6a4b ($p<0.05$) expression). **SF1D)** GABAergic pathway gene expression. In the 24 hpf dnVDRa induced group we see a significant increase in gad67 expression ($p<0.05$). In the 48 hpf dnVDRa induced fish there is a significant increase in gad65 ($p<0.01$) and gad67 ($p<0.05$) expression. In the 72 hpf dnVDRa induced fish there is a significant decrease in gad67 ($p<0.05$) and glsa ($p<0.05$) expression. **SF1E)** Glutaminergic pathway gene expression. In the 24 hpf dnVDRa induced group we see a significant decrease in glula and glulb expression ($p<0.05$) and an increase in slc17a6b ($p<0.05$) expression). In the 48 hpf dnVDRa induced fish there is a significant increase in slc17a7a ($p<0.01$), slc17a6a ($p<0.01$) and slc17a6b ($p<0.05$) expression. In the 72 hpf dnVDRa induced fish there is a significant decrease in glsa ($p<0.05$), glula ($p<0.01$), glulb ($p<0.05$) and slc17a6a ($p<0.05$) expression. **SF1F)** Dopaminergic pathway gene expression. In the 24 hpf dnVDRa induced group we see a significant increase in dat and vmat expression ($p<0.05$) and a decrease in th1 ($p<0.05$), th2 ($p<0.01$) and nr4a2a ($p<0.05$) expression. In the 48 hpf dnVDRa induced fish there is a significant increase in dbh ($p<0.01$), dat ($p<0.05$) and manf ($p<0.001$) expression. In the 72 hpf dnVDRa induced fish there is a significant decrease in dbh ($p<0.01$) and th2 ($p<0.05$) expression and an increase in vmat expression ($p<0.05$).
